# Fortuitous *in vitro* compound degradation produces a tractable hit against *Mycobacterium tuberculosis* dethiobiotin synthetase: a cautionary tale of what goes in, does not always come out

**DOI:** 10.1101/2023.03.26.531482

**Authors:** Wanisa Salaemae, Andrew P. Thompson, Birgit I. Gaiser, Kwang Jun Lee, Michael T. Huxley, Christopher J. Sumby, Steven W. Polyak, Andrew D. Abell, John B. Bruning, Kate L. Wegener

## Abstract

We previously reported potent ligands and inhibitors of *Mycobacterium tuberculosis* dethiobiotin synthetase (*Mt*DTBS), a promising target for antituberculosis drug development (Schumann *et al.,* ACS Chem Biol. 2021, 16, 2339-2347); here the unconventional origin of the fragment compound they were derived from is described for the first time. Compound **1** (9b-hydroxy-6b,7,8,9,9a,9b-hexahydrocyclopenta[3,4]cyclobuta[1,2-c]chromen-6(6a*H*)-one), identified by *in silico* fragment screen, was subsequently shown by surface plasmon resonance to have dose-responsive binding (*K*_D_ 0.6 mM). Clear electron density was revealed in the DAPA substrate binding pocket, when **1** was soaked into *Mt*DTBS crystals, but the density was inconsistent with the structure of **1**. Here we show the lactone of **1** hydrolyses to carboxylic acid **2** under basic conditions, including those of the crystallography soak, with subsequent ring-opening of the component cyclobutane ring to form cyclopentylacetic acid **3**. Crystals soaked directly with authentic **3** produced electron density that matched that of crystals soaked with presumed **1**, confirming the identity of the bound ligand. The synthetic utility of fortuitously formed **3** enabled subsequent compound development into nanomolar inhibitors. Our findings represent an example of chemical modification within drug discovery assays and demonstrate the value of high-resolution structural data in the fragment hit validation process.

**Synopsis:** A molecule flagged in an *in silico* docking screen against *Mt*DTBS, was inadvertently hydrolysed in the crystal conditions used for hit validation. The resulting fragment-sized molecule bound to the DAPA substrate binding pocket of the target enzyme (*Mt*DTBS) with millimolar affinity, as measured by surface plasmon resonance, but was later modified to a highly potent (nanomolar) ligand and promising lead for the development of novel tuberculosis treatments.

**Graphical Abstract:** 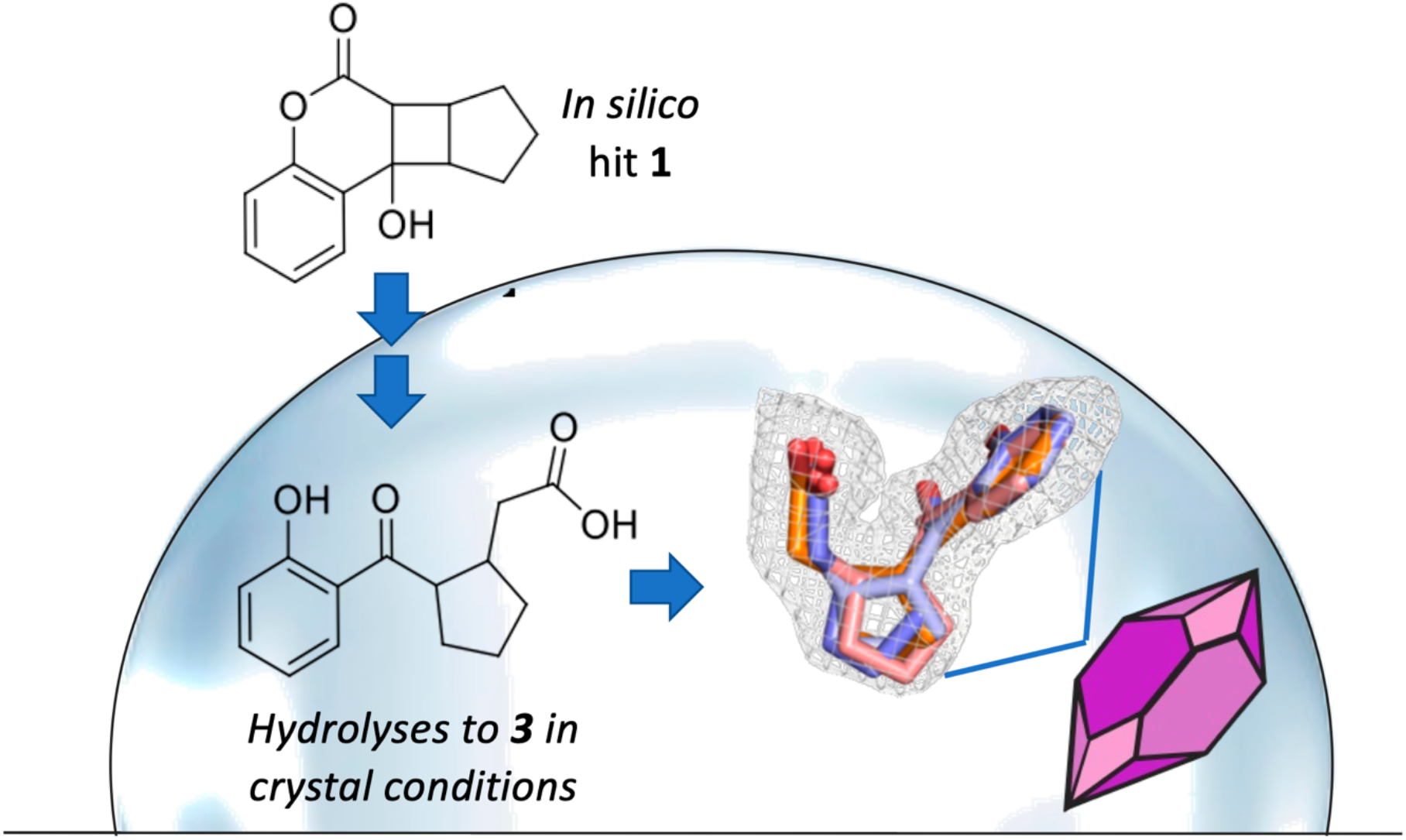

## Introduction

Biotin (vitamin B7) is a cofactor for multiple essential cellular processes including energy metabolism and cell wall maintenance [1–7]. *Mycobacterium tuberculosis* (*Mtb*), the causative pathogen of tuberculosis, relies on biotin biosynthesis as its sole source of this vitamin, as it does not possess the biotin transport proteins found in other bacteria, plants, and animals [1–2]. As a result, targeting enzymes involved in biotin biosynthesis affects the growth of cultured mycobacteria [8–12] as well as their pathogenicity in murine macrophage and zebrafish models of tuberculosis (TB) [13–15], validating biotin biosynthesis as a key metabolic process during both acute and latent stages of the *Mtb* life cycle. The absence of homologous biotin biosynthesis enzymes in humans, suggests targeting this pathway therapeutically may result in a low incidence of off-target effects and is therefore a promising strategy for drug discovery.

Fragment-based drug design (FBDD) has become a favoured alternative to conventional high-throughput drug discovery processes due to its highly efficient sampling of chemical space and ease of subsequent compound modification [16–17]. FBDD has previously been applied to the screening of inhibitors targeting 7,8-diaminopelargonic acid synthase (DAPAS), one of enzymes required for biotin biosynthesis in *Mtb* [18]. The identified inhibitor, an aryl hydrazine, recorded a sub-millimolar inhibition constant (*K*_i_ = 10.4 mM) [18]; however, this is yet to be developed into an anti-TB agent. Dethiobiotin synthetase (DTBS) is another promising antimycobacterial drug target due to its indispensable role in the biotin biosynthesis pathway [2,19]. This protein is well characterised biochemically and structurally [19–20] and was therefore selected for FBDD in the current study.

DTBS functions as an obligate homodimer, with two active sites as revealed by X-ray crystallography [19]. Here, the substrate 7,8-diaminopelargonic acid (DAPA) is converted into dethiobiotin, in a reaction that requires CO2, Mg^2+^, and a nucleoside triphosphate. Each active site is comprised of a DAPA binding site (an enclosed, hydrophobic pocket formed by both dimer partners), and an adjacent nucleoside triphosphate pocket formed solely from one monomer. The latter site is composed of a phosphate binding loop region (P-loop; or Walker A motif) and a nucleoside binding site [19]. DTBS usually requires adenosine triphosphate (ATP) for catalysis, but *M. tuberculosis* DTBS (*Mt*DTBS) can utilise many alternative nucleoside triphosphates (NTPs) in the reaction, with a preference for cytidine triphosphate (CTP) [19–20].

We recently reported the first nanomolar-affinity ligand for *Mt*DTBS derived from a weakly-binding fragment compound [21]. For clarity, this publication focused on the structure-guided development process and details of how the first fragment was identified following the initial *in silico* screen are now reported. We now present the unconventional origin of this fragment, which was produced by hydrolysis of an *in silico* docking hit during crystal soaking experiments designed to confirm the compound binding mode. This result reinforces the potential for compounds to be inadvertently modified during experiments designed to assess their function and demonstrates the value of high-resolution structural data in the hit validation process.

## Results and Discussion

### Initial validation of selected in silico MtDTBS hits by surface plasmon resonance and enzyme inhibition assays

In our previous study, we briefly reported the *in silico* screening of 93,904 fragment compounds from the Zinc database, together with a single concentration SPR follow-up screen of 15 of the highest ranked and available *in silico* hits [21]. Seven of these fragments showed a more than 2-fold increased SPR response over the MgCTP and MgATP controls (B1, B3, B7, B9, B10, B13, and B14 – see Table S1 for structures). We report here the subsequent SPR validation studies of these seven compounds that showed four of the seven, B1, B3, B7, and B9, had dose responsive binding to *Mt*DTBS with *K*_D_ of 0.58 – 1.24 mM (Table S1, Figure S1).

A single dose enzyme inhibition assay was also performed to determine if binding was competitive at the catalytic site. Each of the fifteen originally identified fragments was tested at its maximum soluble concentration, which ranged from 0.18 – 2.39 mM (Figure S2). Only B6, B7, and B8 (at 0.18, 1.38, and 1.05 mM) reduced the *Mt*DTBS activity below 90%, suggesting approximately 10 – 15% inhibition of *Mt*DTBS. The absence of substantial inhibition, particularly for the hits identified as dose-responsive binders by SPR (B1, B3, and B9), may be the result of inadequate solubility as well as the weak binding affinity of the fragments.

### Structural characterisation of hit binding by X-ray diffraction revealed electron density in the MtDTBS active site was incompatible with soaked ligand

Crystals of *Mt*DTBS produced as previously described [19 - 21] were soaked with millimolar concentrations of selected fragments (B1, 45 mM; B7, 5 mM; B9, 30 mM). X-ray diffraction of the soaked crystals showed that in all cases, and consistent with previous observations, DTBS was present as the obligate and catalytically active homo-dimer subunit. Electron density corresponding to a soaked ligand in the active site was only observed with compound B9.

Crystals soaked with fragment B9, re-designated here for simplicity as compound **1** (9b-hydroxy-6b,7,8,9,9a,9b-hexahydrocyclopenta[3,4]cyclobuta[1,2-c]chromen-6(6aH)-one; 30 mM final concentration), were shown to have additional electron density in the DAPA pocket, between two bound sulfate ions. This density differed from that expected for compound **1** based on the Zinc database (Figure 1A), and did not match that expected for any of the crystallographic buffer components present, i.e. sulfate, glycerol and DMSO.

**Figure 1.**
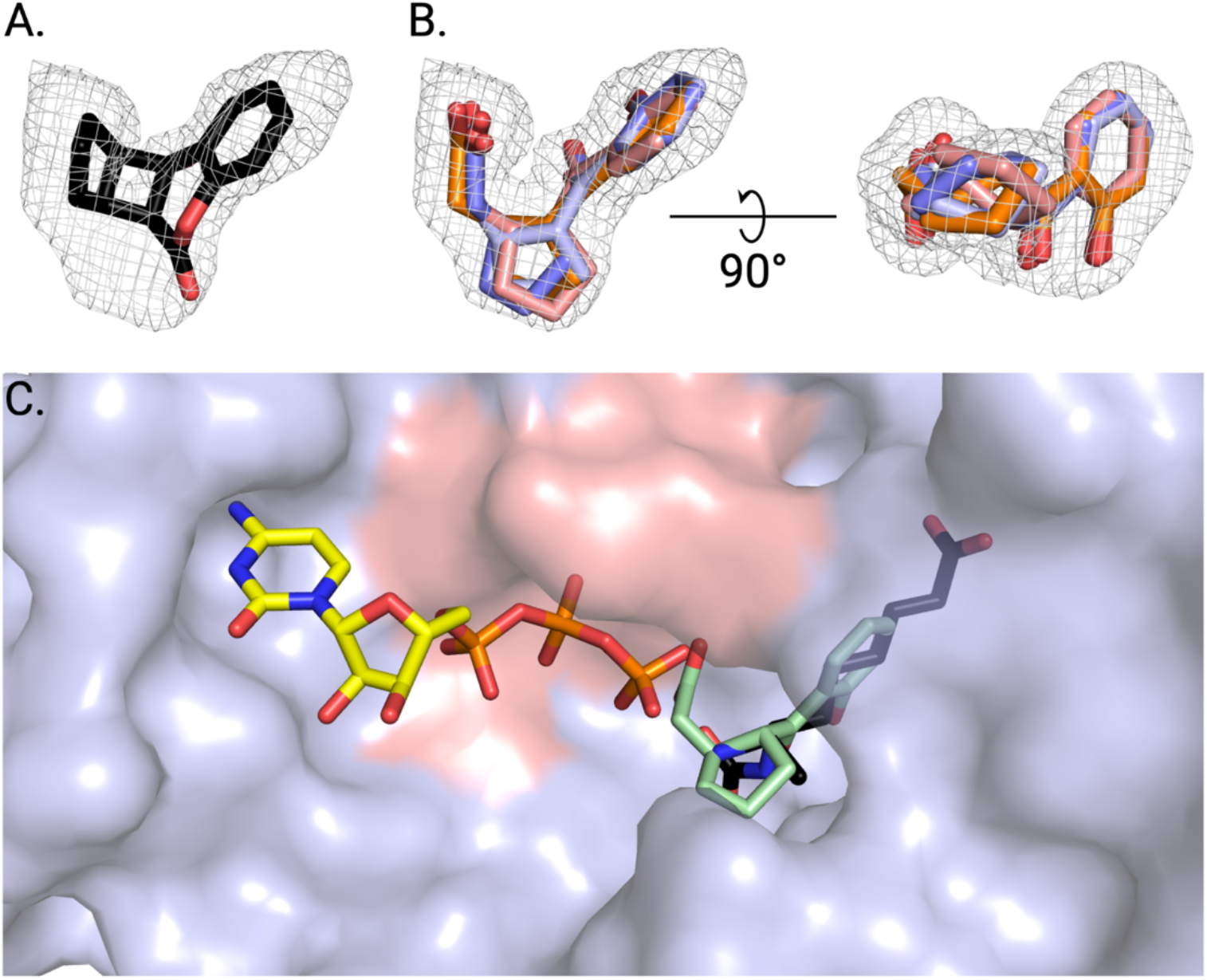
Electron density from an MtDTBS crystal soaked with compound **1**. **A.** View of compound **1** (black sticks) within the Polder electron density (3s) observed within the active site of MtDTBS. This molecule fits poorly into the density. **B.** Two perspectives of each diastereomer of **3** (RR: light blue, SS: dark blue, SR: pink, RS: orange) fit into the Polder electron density (3s) observed in the MtDTBS active site. Each of these molecules is visually consistent with the electron density. Minimal structural differences between diastereomers were observed, and the preferred stereochemistry could not be definitively identified solely by crystallography. **C.** Compound **3** overlayed with the natural substrates of MtDTBS: CTP (yellow sticks) and DAPA (black sticks). **3** binds in the DAPA pocket.

It is noted here, that lactone **1** was described in the Zinc database as the *cis,cis,cis-* stereoisomer (Figure 2). However, X-ray diffraction of a small crystal that had grown in the stock solution of **1** allowed the molecule to be reassigned the *trans,trans,cis* relative configuration shown in Figure 2 (Table S2). It was later noted that the supplier linked to this compound by the Zinc database (Specs Compound Handling BV) did not specify the configuration around the cyclobutane ring. Despite this ambiguity over the starting relative configuration, both the *trans,trans,cis-* and *cis,cis,cis*-stereoisomers were clearly incompatible with the *Mt*DTBS ligand electron density.

**Figure 2.**
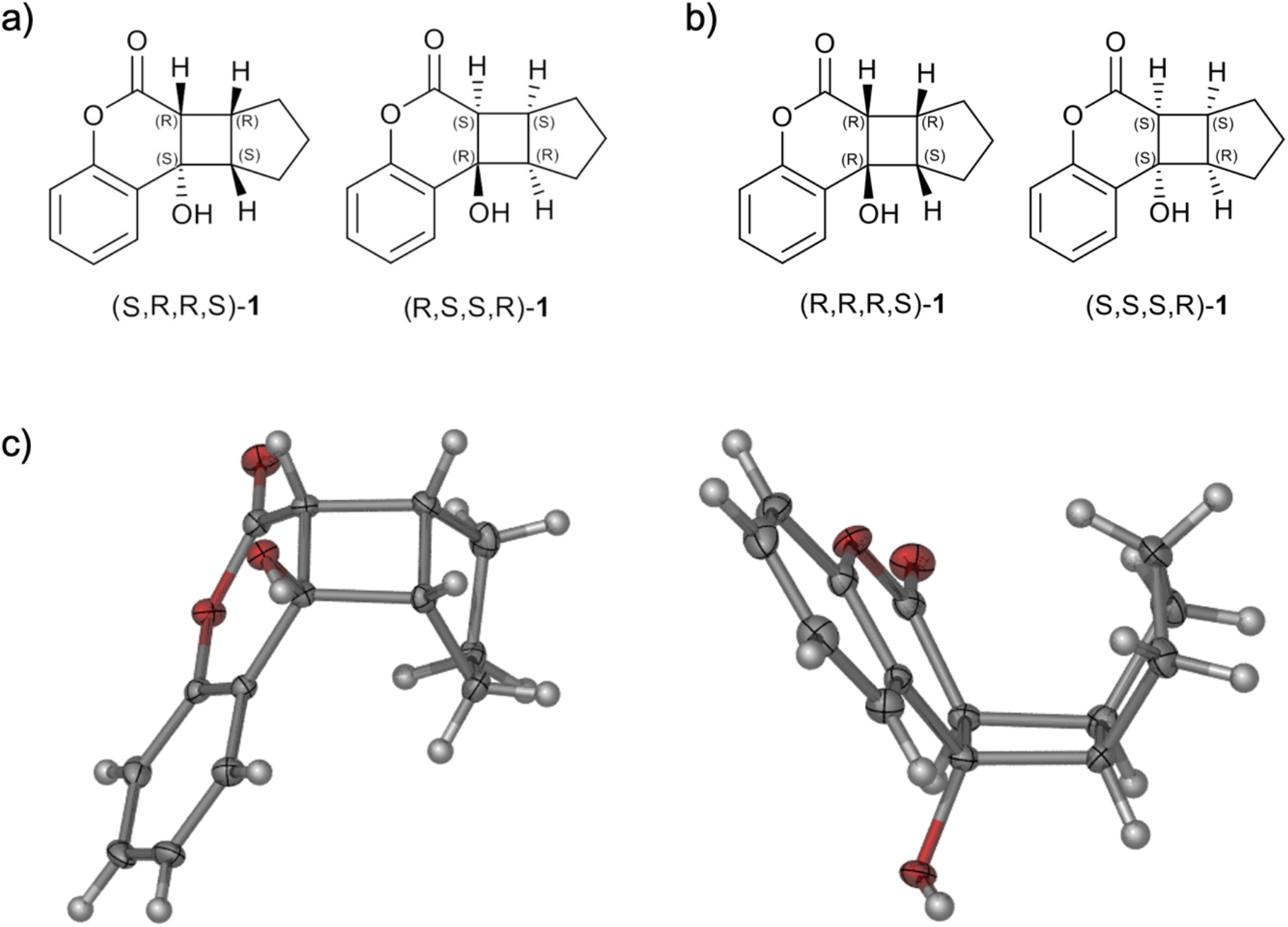
The enantiomers and the relative stereochemistry of lactone **1** a) as described in the Zinc database, and b) as determined by X-ray crystallography. Panel c) shows the three-dimensional view of **1**. Both enantiomers were observed in the asymmetric unit of the crystal. Displacement ellipsoids are shown at the 50% probability level.

It was hypothesised that the lactone of **1**, fused to a strained cyclobutane ring, would be susceptible to hydrolysis to intermediate cyclobutane **2** with subsequent ring opening giving carboxylic acid **3** (Scheme 1). Compound **3** was more visually consistent with the observed electron density present in the active site of *Mt*DTBS and modelling of this molecule in the 2Fo-Fc electron density resulted in favourable COOT density fit scores and PHENIX real-space correlation coefficients (>1.09, >0.938 respectively). Modelling was not sufficient to conclusively determine either the relative or absolute configuration of the bound compound. Each stereoisomer of structure **3** could be manipulated to fit within the density (Figure 1B); however, upon refinement of one of the *trans*-**3** exhibited the highest ligand occupancy (RS: 0.51, SR: 0.13, RR: 0.32, SS: 0.34, RS and SR are *trans,* SS and RR are *cis*) and the most preferable real-space refinement statistics (Table S3).

**Scheme 1.**
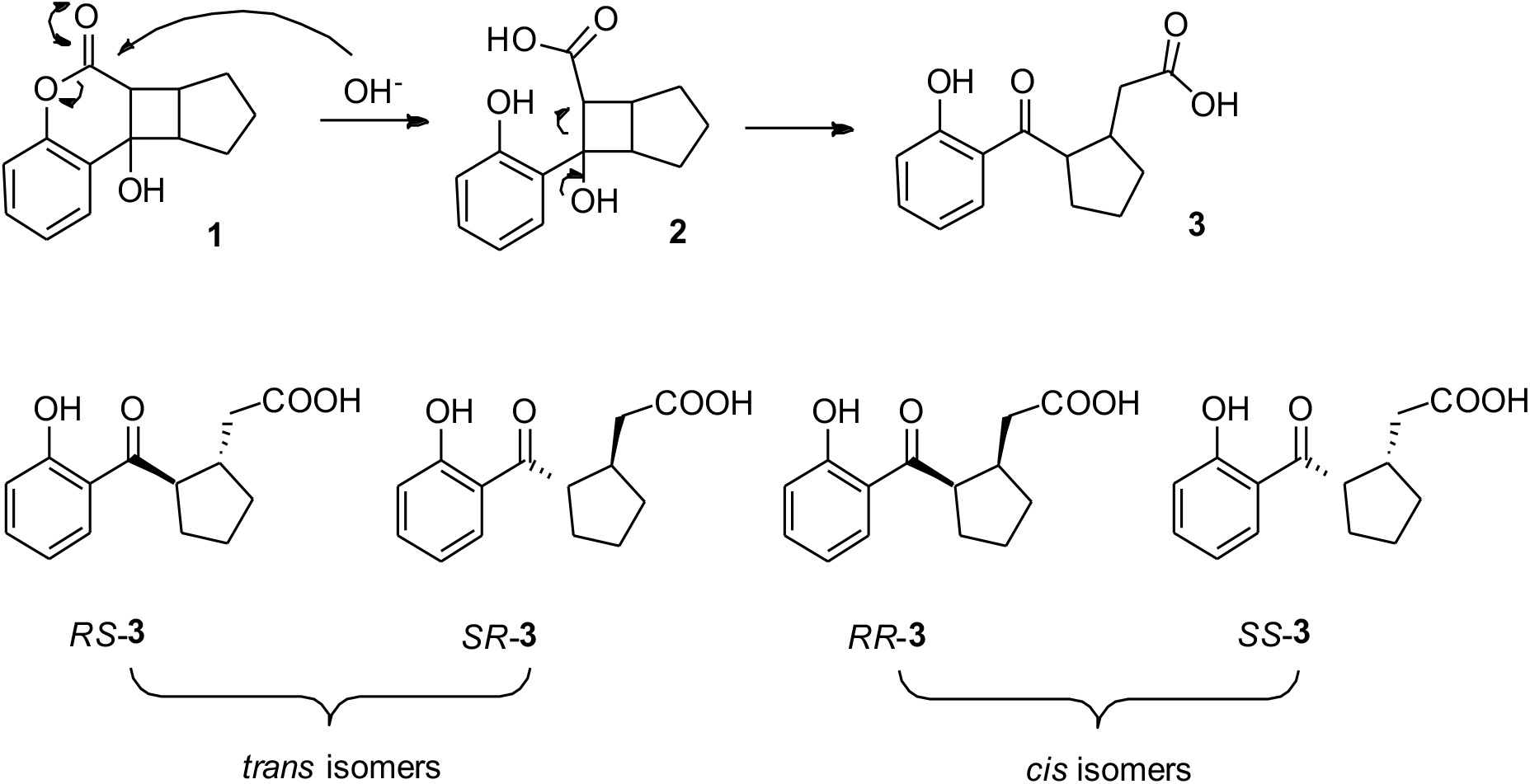
Proposed degradation mechanism of lactone **1** to carboxylic acid **3**, via cyclobutane **2** in the crystallization conditions. Degradation was hypothesized to occur via hydrolysis of the lactone and collapse of the four-membered ring. This was accompanied by the generation of two new stereocenters, leading to four possible diastereomers (RS, SR, RR, and SS).

### Basic conditions in crystal soaking induced the hydrolysis of compound 1

To test the hypothesis that the density observed in the *Mt*DTBS active site was due to a hydrolysis product, a combination of small molecule X-ray crystallography, NMR, TLC, and MS analysis was performed. Firstly, deliberate hydrolysis of **1** using classical basic hydrolysis conditions [22], produced carboxylic acid **3** (Scheme 2).

**Scheme 2.**
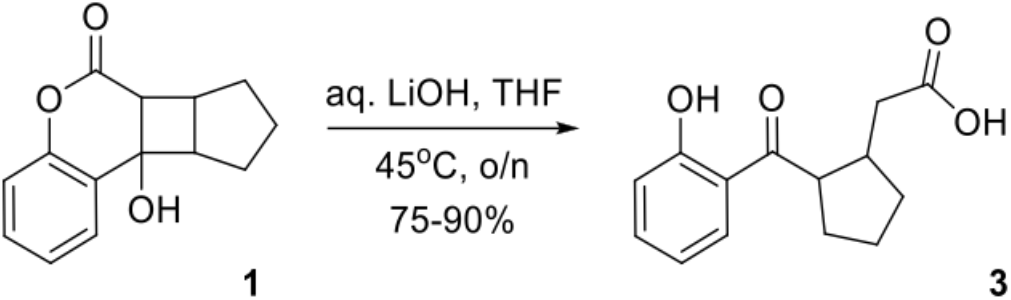
*Induced hydrolysis of lactone **1** to carboxylic acid **3** under basic condition. The presence of **3** was confirmed by NMR and MS analysis (see data in* Compound Synthesis, *Methods section).*

To investigate whether compound **1** had hydrolysed in the original stock solution, or under the crystal soaking conditions, we examined the DMSO stock solution of **1** used for these experiments, for signs of hydrolysis. More than one year since its original preparation, the stock solution was found to contain a 1:1 mixture of both **1** and **3**, as determined by NMR analysis (Figure S3).

Next, the effect of the slightly basic crystal soaking conditions (pH 8) on compound **1** was investigated. A small amount of **1** was added to the crystal soaking buffer (1.4 M NH4SO4, 0.1 M Tris pH 8, 10% glycerol), and the reaction mixture was monitored by TLC over the course of a week. The first traces of **3** were detected after 2 days, with complete conversion after 6 days (data not shown). This contrasted with the forced hydrolysis of **1**, that was complete within 24 hours.

Together, these results showed that **1** only slowly degraded to **3** in neutral aqueous DMSO, but that degradation was significantly hastened by increasingly basic aqueous conditions. Compound **3** observed in the crystal structure was likely predominantly produced during the 8-day crystal soak.

### Compound 3 is a trans-stereoisomer

Based on the proposed degradation mechanism (Scheme 1) and the *trans,trans,cis-* stereochemistry of **1**, it is conceivable that the substituents on the cyclopentane ring in **3** also have a *cis*-relative configuration (see Scheme 1). However, the proton α to the ketone of **3** is expected to be base labile and hence could potentially epimerise. Unfortunately, no small molecule crystal of **3** could be obtained and NOESY NMR experiments were inconclusive. The relative stereochemistry of **3** was instead indirectly determined with the help of compound **4** (Figure 3). This ethyl ester was synthesized as part of efforts to prepare analogues of compound **3** and surprisingly consisted of two separable diastereoisomers as shown in Figure 4. A NOESY spectrum was recorded for each diastereomer, and it was concluded that the *cis*-isomer was that which showed a correlation between the two CH protons on the cyclopentane ring, whereas the *trans*-isomer showed a correlation between the ketone α-proton and the methylene group attached to the cyclopentane ring (Figure 3). Notably, the α-proton of *trans-**4*** was shifted significantly upfield relative to that of *cis-**4*** (δ_cis_ ≈ 3.9ppm; δ_trans_ ≈ 3.5 ppm).

**Figure 3.**
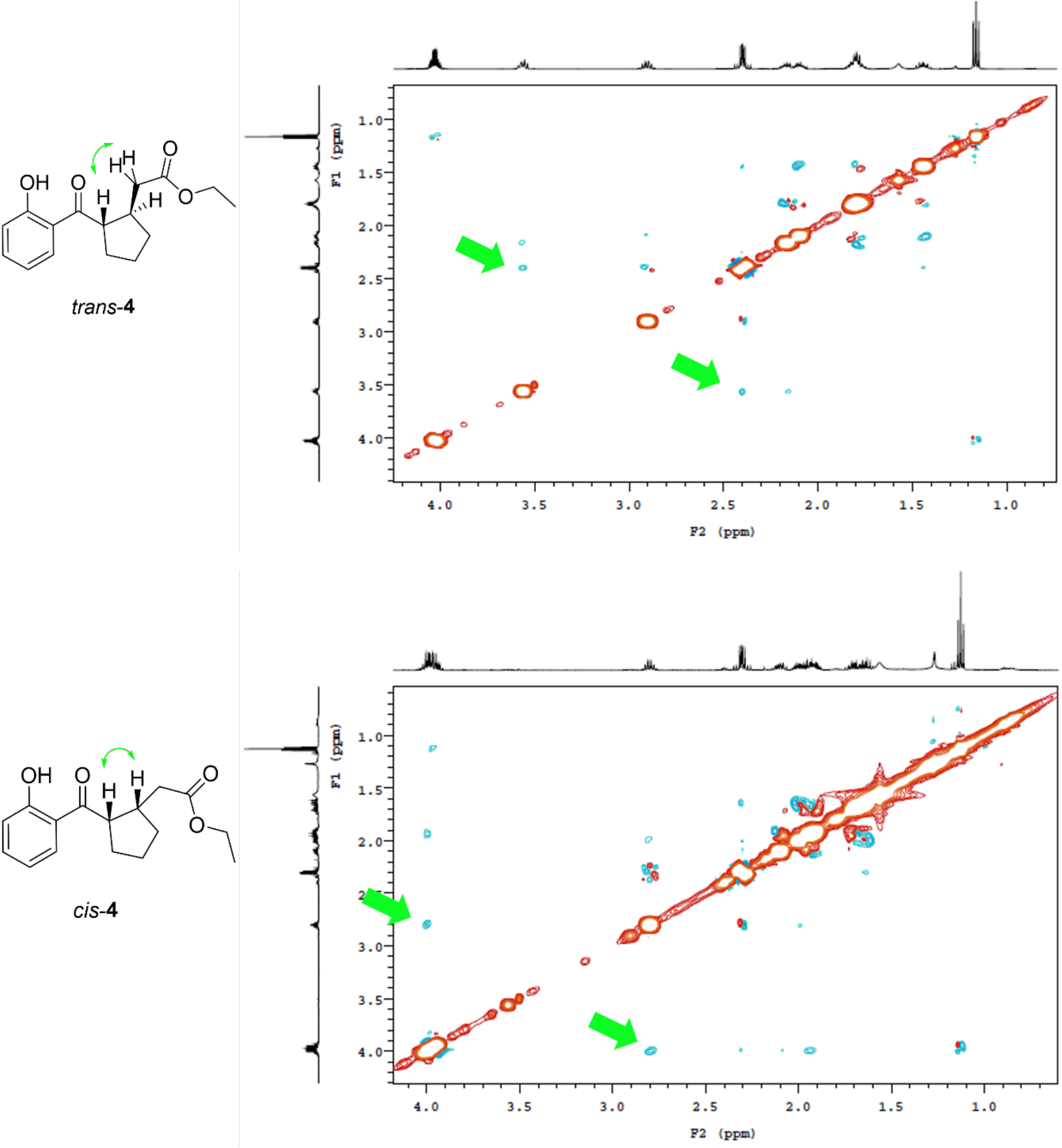
Structure of trans and cis isomers of 4 (left) and corresponding NOESY spectra (right). Green arrows indicate the key correlations found in the NOESY experiments. Stereochemical configurations are relative.

Comparison of the ^1^H NMR spectra of both diastereoisomers of **4** with that of **3**, as produced by forced hydrolysis, led to the conclusion that **3** obtained in this way is in its *trans*conformation (Table S4). As noted above, the ligand density observed in the *Mt*DTBS cocrystal structure was also consistent with the *trans* conformation (Figure 1). Together the X-ray diffraction of the *Mt*DTBS co-crystal and NMR data confirm the identity of the hydrolysed product as *trans*-**3**. As an additional test, *Mt*DTBS crystals soaked with **3** produced by deliberate hydrolysis (PDBID 6NMZ [21]) had electron density in the active site that closely matched that obtained after soaking with **1** (PDBID: 6NLZ, shown in Figure 1) and *trans* isomers modelled into the density from each crystal were well aligned (RMSD align = 0.181 - 0.278, Figure 4).

**Figure 4.**
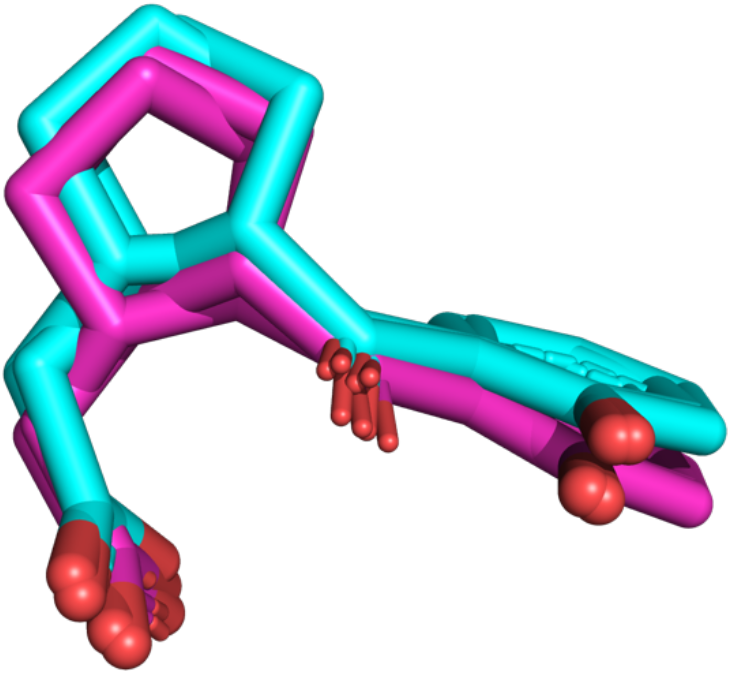
An overlay of **3** from the PDBIDs 6NLZ (cyan, soaked directly with **1** – compound **3** forms during the soaking period) and 6NMZ (magenta, soaked with **3** formed by prior hydrolysis of **1**). This overlay depicts that the intentionally produced **3** exhibits the same binding pose as the original observation of the molecule.

## Conclusions

In this report, we describe the serendipitous discovery of a fragment molecule that binds to *Mt*DTBS, a promising anti-tuberculosis drug target. Compound **1**, which was initially identified by *in silico* screening, degraded when added to the crystal soaking condition, resulting in **3**, which bound to the DAPA substrate pocket. Compound **3** was later determined to bind only weakly to *Mt*DTBS with a *K*_D_ of 3.4 mM [21]. This was consistent with the requirement for high concentrations for the crystal soaking, as electron density for **3** was not observed in soaks with concentrations below 10 mM. Meanwhile, here the *K*_D_ of compound **1** was measured as 0.6 mM (Figure S1), indicating the original compound had stronger binding affinity than hydrolysis product **3**. However, weak *K*_D_S in the high micromolar-millimolar range can be hard to measure accurately and are associated with large errors. In addition, different methods were used to immobilise the *Mt*DTBS for these two experiments (biotinylated-DTBS coupling to streptavidin for **3** [21] and amine coupling for **1**, reported here), which may influence the results. Nevertheless, these results show that both **1** and **3** bind to *Mt*DTBS.

Although compound **1** bound to *Mt*DTBS with ostensibly higher affinity than compound **3**, it is a more complex and unstable molecule as revealed here. This instability prevents access to high resolution X-ray structural data at least under the current crystallization conditions. On the other hand, fragment **3** with its carboxyl group has far greater synthetic utility than **1**, is chemically more stable due to the absence of the lactone, and is amenable to high resolution structural characterisation. As such, relatively simple but rational chemical modifications to compound **3**, including addition of a tetrazole substituent on the aromatic ring, removal of the phenol group and inclusion of a further carboxylic acid group, improved the affinity over 5 orders of magnitude to a final *K_D_* of 57 nM [21]. The finding in this study constitutes an example of chemical tractability trumping hit affinity, in the prioritisation of hits.

The need to verify compound integrity during storage and transport has been known for some time [23–24] but even regular quality control of a fragment library may not have been sufficient in the case of compound **1**, since the degradation occurred mainly within the crystallization solution, rather than the stock. This occurrence highlights that library quality control should also be performed in media used in experiments, over an experimentally relevant time-frame. This could be done concurrently with compound screening in the case of ligand detected NMR [25–26]. However, other common screening techniques such as SPR and thermal shift assays would not be able to detect compound degradation such as that presented here. Parallel analysis of promising hits under the same screening conditions, for example by NMR or LC/MS, may be beneficial in these cases.

This report also highlights the importance of high-resolution structural data at early stages of fragment-based projects. In the absence of such data, optimisation of compound **1** would have been highly challenging. X-ray crystallography provided invaluable data, not only for observing compound degradation, but also allowing the structure-based chemical development of **3** to achieve a >10^7^-fold increase in affinity for *Mt*DTBS [21]. This presents a strong argument for the inclusion of X-ray crystallographic data at early stages in the FBDD process.

## Methods

### Surface plasmon resonance analysis

SPR analysis was performed using a BIAcore T100 instrument. The preparation of *Mt*DTBS was as previously described [19]. Proteins were diluted to 0.2 mg mL^-1^ in 0.01 M NaOAc buffer (pH 5.2) and immobilized to the surface of a CM5 sensor chip using a constant flow rate of 5 μl min^-1^ and contact time of 420 s to reach approximately 10,000 final response units (RU). Flow cell 1 was left blank (no protein) to correct for bulk refractive index changes, flow cell 2 was used to immobilized *Mt*DTBS. 2.5% Dimethyl sulfoxide (DMSO) was added to the binding buffer (10 mM HEPES pH 7.4, 150 mM NaCl, and 0.005% (v/v) surfactant P20) to alleviate solubility problems. To correct for small dilution errors during the preparation of compounds, a solvent correction was performed using standards containing 2% to 3.3% (v/v) DMSO. Hits previously identified by SPR as having ≥ 2-fold increase in SPR response above the MgCTP and MgATP controls at a single concentration of compound (1 mM) [21] were subjected to binding analysis using varying concentrations (0.1 – 3 mM) to determine the binding affinity (*K*_D_). The *K*_D_ was determined using a steady-state kinetic model with BIAcore T100 evaluation software (GE Healthcare).

### Enzyme inhibition assay

The enzyme activity of *Mt*DTBS was measured using a Fluorescence Polarization (FP)-based detection assay was previously described [19]. In the inhibition assay, test compound dissolved in 1% DMSO was included in the enzyme activity assay containing *Mt*DTBS (1.5 μM) and saturating concentrations of DAPA (0.1 mM) and MgATP (0.3 mM). The synthesized DTB product was calculated in comparison with the standard DTB curve. The control reaction was performed in the absence of test compound and this amount of synthesized DTB product from the reaction was considered as the reflection of 100% enzyme activity. The inhibitory activity of compounds was determined by the percentage of the remaining enzyme activity.

### X-ray crystallography of MtDTBS complexes

*Mt*DTBS crystals were grown in 1.2 – 1.7 M ammonium sulfate, 0.1 M Tris pH 8, 10-15% glycerol via hanging drop vapour diffusion at 16°C as previously described [20]. *Mt*DTBS crystals were soaked with compound **1** 9b-hydroxy-6b,7,8,9,9a,9b-hexahydrocyclopenta[3,4]cyclobuta[1,2-c]chromen-6(6aH)-one (30 mM final concentration; 10% DMSO) for 8 days. Synthesized-**3**-*Mt*DTBS crystals were soaked with **3** (20 mM final concentration; 10% DMSO) for 2 days. Soaked crystals were subsequently flash cooled in liquid nitrogen and subjected to X-ray diffraction on the MX-1 Beamline at the Australian Synchrotron. Structures were indexed, scaled, and merged in either iMosflm and AIMLESS (CCP4) in accordance with CC1/2 cut-off values. The phase problem was solved via molecular replacement (PhaserMR or DIMPLE) using in-house *Mt*DTBS search models. All crystal structures were solved in the P212121 space group, with identical overall fold to previous structures [19–21]. Structures were refined in Coot and Phenix.refine.

### Small molecule X-ray crystallography

A small crystal of B9/Compound **1** formed over an indeterminate time in the DMSO stock solution, prepared from commercially obtained compound (Specs Compound Handling, BV). Data was collected on an Xcalibur diffractometer fitted with an Eos detector on a crystal mounted on a nylon loop at 150(2) K. Using Olex2 [27], the structure was solved with the Direct Methods using the SHELXS [28] structure solution program and refined with the SHELXL [29] refinement package using Least Squares minimisation. Crystal and structure refinement data are provided in Table S2.

### Thin-Layer Chromatography (TLC)

Pre-coated silica gel 60 F254 plates purchased from Merck were used for TLC. EtOAc, petroleum benzine, DCM, MeOH, and mixtures thereof were used as eluents (see compound synthesis section for specific individual conditions). Visualization of the compounds was achieved with UV light (254 nm). The denoted retention factors *(Rf)* were rounded to the nearest 0.05.

### High Performance Liquid Chromatography (HPLC)

HPLC retention times (*R*_T_) are reported in minutes (min) and were determined by different methods, given in parenthesis. HPLCs were recorded on an Agilent 1260 Infinity HPLC and detector (detection at 254 nm) with a phenomenex Luna 5 μm C18 (2) 100 Å (250 X 4.60 mm) connected using mobile phase (MP) A: 0.1% TFA in Milli-Q water, MP B: 0.08% TFA in acetonitrile. For Method A, flowrate: 1 mL/min. Gradient: 0% B over 5 min, 0-100% B over 15 min, and 100% B over 2 min. For Method B, flowrate: 1 mL/min. Gradient: 60% B over 5 min, 60-100% B over 15 min, and 100% B over 3 min.

### Nuclear Magnetic Resonance Spectroscopy (NMR)

NMR experiments were performed on a Varian Inova 500 MHz or 600 MHz instrument. The obtained spectra were analysed using MestReNova 11.0 software typically using Whittaker smoother baseline correction. Chemical shifts are reported in ppm (δ) with reference to the deuterated solvent used [30]. Coupling constants *(J)* are reported in Hertz (Hz) and rounded to the nearest 0.5 Hz. Multiplet patterns are designated the following abbreviations or combinations thereof: m (multiplet), s (singlet), d (doublet), t (triplet), q (quartet). Signal assignment was made from unambiguous chemical shifts or coupling constants and COSY, HSQC and NOESY experiments.

### High Resolution Mass Spectrometry (HRMS)

High resolution mass spectra were recorded on Agilent 6230 time of flight (TOF) liquid chromatography mass spectra instrument.

### Compound synthesis

#### 2-(2-(2-hydroxybenzoyl)cyclopentyl)acetic acid (3)

Compound **3** - 2-(2-(2-hydroxybenzoyl)cyclopentyl)acetic acid - was produced by classical hydrolysis of 9b-hydroxy-6b,7,8,9,9a,9b-hexahydrocyclopenta[3,4]cyclobuta[1,2-c]chromen-6(6aH)-one (**1**) under basic conditions. Compound **1**, obtained from Specs Compound Handling BV (0.50 g, 2.17 mmol), was dissolved in THF (20 mL) and 1 N aq. LiOH (9 mL) was added. The reaction mixture was stirred at 45 °C for 18 hr before it was acidified with 2 N aq. HCl (ca. 50 mL) and the product was extracted into EtOAc (4 × ca. 50 mL). The combined organic layers were washed with brine (ca. 50 mL), dried over MgSO4 and concentrated *in vacuo.* Column chromatography (8% MeOH in DCM, v/v), following standard procedures using silica gel 60 (40-63 μm mesh), gave the product as a pale-yellow oil (0.48 g, 90%). **TLC** *R*_f_ = 0.35 (5% MeOH in DCM, v/v); **HPLC** *R*_T_ = 18.59 min (Method A); **^1^H NMR** (500 MHz; CDCl3) δ 12.45 (s, 1H, OH), 7.76 (dd, *J* = 8.0, 1.5 Hz, 1H, ArH), 7.49 – 7.43 (m, 1H, ArH), 6.98 (d, *J* = 8.5 Hz, 1H, ArH), 6.90 (t, *J* = 7.6 Hz, 1H, ArH), 3.52 (q, *J* = 8.5, 8.0 Hz, 1H, (C=O)CH), 2.89 (dq, *J* = 15.5, 8.0 Hz, 1H, (C=O)CHC*H*), 2.47 (dd, *J* = 15.3, 6.4 Hz, 1H, (COOH)C*H_A_*H_B_), 2.37 (dd, *J* = 15.3, 7.9 Hz, 1H, (COOH)CH_A_*H_B_*), [2.23 – 2.06 (m, 2H), 1.83 – 1.71 (m, 3H), 1.51 – 1.37 (m, 1H) (3×CH2)]; **^13^C NMR** (126 MHz; CDCl3) δ 208.4 (C=O), 178.0 (COOH), 163.1 (*C_Ar_*C=O), 136.4 (C_Ar_H), 130.3 (C_Ar_H), 119.4 (C_Ar_OH), 119.0 (C_Ar_H), 118.7 (C_Ar_H), 51.5 ((C=O)*C*H), 38.6 (*C*H2COOH), 38.6 ((C=O)CH*C*H), 32.6, 32.3, 24.9 (3×CH2); **HRMS** *m/z* (+ESI) found: 271.0943 [M+Na]^+^; C_14_H_16_O_4_Na^+^ requires *M,* 271.0941.

#### Ethyl 2-(2-(2-hydroxybenzoyl)cyclopentyl)acetate (4)

9b-hydroxy-6b,7,8,9,9a,9b-hexahydrocyclopenta[3,4]cyclobuta[1,2-c]chromen-6(6aH)-one (**1**) (40 mg, 0.17 mmol) was dissolved in EtOH (2 mL), and methyl-*L*-serinate hydrochloride (27 mg, 0.17 mmol) and NaHCO3 (16 mg, 0.19 mmol) were dissolved in EtOH/H2O (1:1, 2 mL), and added [31]. The solution was stirred at ambient temperature for 18 hr, before it was heated to reflux for 5 hr until full conversion of the starting material. After cooling to r.t. the reaction mixture was acidified with aq. 2 N HCl (ca. 10 mL). The product was extracted into EtOAc (3 × ca. 20 mL), and the combined organic layers were washed with brine (ca. 30 mL), dried over MgSO4 and concentrated *in vacuo*. The product was purified by column chromatograpy (10% EtOAc in petroleum benzine (v/v)) to give **4** as a colourless oil and a mixture of diastereoisomers (21 mg, 34%). The isomers were separated by semi-preparative HPLC to give *cis*-**4** (7 mg, 11%, 86% de) and *trans*-**4** (14 mg, 23%, 94% de).

##### *cis*-isomer

**TLC** *R*f = 0.40 (10% EtOAc in petroleum benzine, v/v); **HPLC** *R*_T_ = 14.01 min (Method B); **^1^H NMR** (500 MHz; CDCl3) *δ* 12.54 (s, 1H, OH), 7.81 (dd, *J* = 8.0, 1.5 Hz, 1H, ArH), 7.49 – 7.43 (m, 1H, ArH), 6.98 (t, *J* = 6.0 Hz, 1H, ArH), 6.92 – 6.87 (m, 1H, ArH), 4.06 – 3.87 (m, 3H, OCH2 + CHC=O), 2.83 – 2.73 (m, 1H, C*H*CHC=O), 2.34 – 2.22 (m, 2H, CH2C=O), [2.13 – 2.04 (m, 1H), 2.03 – 1.84 (m, 3H), 1.73 – 1.59 (m, 2H) (3xCH2)], 1.14 – 1.08 (m, 3H, CH3); **^13^C NMR** (126 MHz; CDCl3) *δ* 209.1 (Ar*C*=O), 172.9 (CH2*C*=O), 163.0 (C_Ar_OH), 136.5, 130.3 (2xC_Ar_H), 120.0 (*C_Ar_*C), 119.0, 118.8 (2xC_Ar_H), 60.5 (OCH2), 47.9 (*C*HC=O), 40.5 (*C*HCHC=O), 35.9 (*C*H2C=O), 32.5, 29.1, 23.9 (3xCH2), 14.2 (CH3); **HRMS** *m/z* could not be detected.

##### *trans*-isomer

**TLC** *R*f = 0.40 (10% EtOAc in petroleum benzine, v/v); **HPLC** *R*_T_ = 13.51 min (Method B); **IR** *ν*_max_(neat)/cm^-1^ 2962, 1731 (C=O), 1632 (C=O), 1446, 1259, 1014, 793, 756; **^1^H NMR** (500 MHz; CDCl3) *δ* 12.50 (s, 1H, OH), 7.77 (dd, *J* = 8.0, 1.5 Hz, 1H, ArH), 7.50 – 7.43 (m, 1H, ArH), 6.98 (dd, *J* = 8.4, 1.0 Hz, 1H, ArH), 6.93 – 6.87 (m, 1H, ArH), 4.08 – 3.96 (m, 2H, OCH2), 3.58 – 3.51 (m, 1H, CHC=O), 2.95 – 2.83 (m, 1H, C*H*CHC=O), 2.45 – 2.33 (m, 2H, CH2C=O), [2.21 – 2.01 (m, 2H), 1.86 – 1.69 (m, 3H), 1.48 – 1.38 (m, 1H) (3xCH2)], 1.15 (t, *J* = 7.0 Hz, 3H, CH3); **^13^C NMR** (126 MHz; CDCl3) *δ* 208.7 (Ar*C*=O), 172.6 (CH2*C*=O), 163.1 (C_Ar_OH), 136.3, 130.3 (2xCArH), 119.5 (*C_Ar_*C), 119.0, 118.7 (2xC_Ar_H), 60.5 (OCH2), 51.5 (*C*HC=O), 39.4 (*C*H2C=O), 39.0 (*C*HCHC=O), 32.9, 32.4, 25.0 (3xCH2), 14.2 (CH3); **HRMS** *m/z* could not be detected.

## Supporting Information

Table S1-S2 and Figure S1-3 are provided in Salaemae_et_al_supp_info.pdf. Atomic coordinates and structure factors of *Mt*DTBS soaked with compound **1** (ligand density corresponds to **3**) have been deposited in the Protein Data Bank, Research Collaboratory for Structural Bioinformatics, Rutgers University, New Brunswick, NJ, with PDBID: 6NLZ. The structural data obtained for pure compound **1** were deposited in the Cambridge Crystallographic Data Centre, with CCDC reference number 2237689.

## Acknowledgements

This research was supported by a Channel 7 Children’s Research Foundation Grant (181614) awarded to K.L.W., A.A., J.B.B., and S.W.P., and a National Science, Research and Innovation Fund (NSRF) and Prince of Songkla University (Grant No. MED6505096i) awarded to W.S. This research was undertaken in part using the MX1 beamlines at the Australian Synchrotron, part of ANSTO, and made use of the Australian Cancer Research Foundation (ACRF) detector.

## Notes

### Competing Interest Statement

The authors have declared no competing interest.

## References

1. Salaemae W, Azhar A, Booker GW, Polyak SW. Biotin biosynthesis in *Mycobacterium tuberculosis:* physiology, biochemistry and molecular intervention. Protein Cell. 2011 Sep;2(9):691–5. Available from: DOI: 10.1007/s13238-011-1100-8

2. Salaemae W, Booker GW, Polyak SW. The role of biotin in bacterial physiology and virulence: a novel antibiotic target for *Mycobacterium tuberculosis*. Microbiol Spectr. 2016 Apr;4(2). Available from: DOI: 10.1128/microbiolspec.VMBF-0008-2015.

3. Park S, Klotzsche M, Wilson D, Boshoff H, Eoh H, Manjunatha U, Blumenthal A, Rhee K, Barry III C, Aldrich C, Ehrt S, Schappinger D. Evaluating the sensitivity of *Mycobacterium tuberculosis* to biotin deprivation using regulated gene expression. PLoS Pathog. 2011 7:1–10.

4. Eisenreich W, Dandekar T, Heesemann J, Goebel W. Carbon metabolism of intracellular bacterial pathogens and possible links to virulence. Nat Rev Microbiol. 2010 8:401–412.

5. Takayama K, Wang C, Besra G. Pathway to synthesis and processing of mycolic acids in *Mycobacterium tuberculosis*. Clin Microbiol Rev. 2005 18:81–101.

6. Tong L. Structure and function of biotin-dependent carboxylases. Cell Mol Life Sci. 2013 70:863–891.

7. Baez-Saldana A, Zendejas-Ruiz I, Revilla-Monsalve C, Islas-Andrade S, Cardenas A, Rojas-Ochoa A, Vilches A, Fernandez-Mejia C. Effects of biotin on pyruvate carboxylase, acetyl-CoA carboxylase, propionyl-CoA carboxylase, and markers for glucose and lipid homeostasis in type 2 diabetic patients and nondiabetic subjects. Am J Clin Nutr. 2004 79:238–243.

8. Mann S, Marquet A, Ploux O. Inhibition of 7,8-diaminopelargonic acid aminotransferase by amiclenomycin and analogues. Biochem Soc Trans. 2005 33:802–805.

9. Singh G, Singh G, Jadeja D, Kaur J. Lipid hydrolizing enzymes in virulence: *Mycobacterium tuberculosis* as a model system. Crit Rev Microbiol. 2010 36:259–269.

10. Deb C, Lee C, Dubey V, Daniel J, Abomoelak B, Sirakova T, Pawar S, Rogers L, Kolattukudy P. A novel *in vitro* multiple-stress dormancy model for *Mycobacterium tuberculosis* generates a lipid-loaded, drug-tolerant, dormant pathogen. PLoS One. 2009 4:1–15.

11. Keer J, Smeulders M, Gray K, Williams H. Mutants of *Mycobacterium smegmatis* impaired in stationary-phase survival. Microbiology. 2000 146:2209–2217.

12. Sassetti C, Boyd D, Rubin E. Comprehensive identification of conditionally essential genes in mycobacteria. Proc Natl Acad Sci. 2001 98:12712–12717.

13. Yu J, Niu C, Wang D, Li M, Teo W, Sun G, Wang J, Liu J, Gao Q. MMAR_2770, a new enzyme involved in biotin biosynthesis, is essential for the growth of *Mycobacterium marinum* in macrophages and zebrafish. Microbes Infect. 2011 13:33–41.

14. Rengarajan J, Bloom B, Rubin E. Genome-wide requirements for *Mycobacterium tuberculosis* adaptation and survival in macrophages. PNAS. 2005 102:8327–8332.

15. Sassetti C, Rubin E. Genetic requirements for mycobacterial survival during infection. Proc Natl Acad Sci. 2003 100:12989–12994.

16. Erlanson DA, Fesik SW, Hubbard RE, Jahnke W, Jhoti H. Twenty years on: the impact of fragments on drug discovery. Nat Rev Drug Discov. 2016 15:605–619.

17. Lamoree B, Hubbard RE. Using fragment-based approaches to discover new antibiotics. SLAS Discov 2018 23:495–510.

18. Dai R, Wilson D, Geders T, Aldrich C, Finzel D. Inhibition of *Mycobacterium tuberculosis* transaminase BioA by aryl hydrazine and hydrazides. Chembiochem. 2014 15:575–586.

19. Salaemae W, Yap MY, Wegener KL, Booker GW, Wilce MC, Polyak SW. Nucleotide triphosphate promiscuity in *Mycobacterium tuberculosis* dethiobiotin synthetase. Tuberculosis (Edinb). 2015 May;95(3):259–266. Available from: DOI: 10.1016/j.tube.2015.02.046.

20. Thompson AP, Salaemae W, Pederick JL, Abell AD, Booker GW, Bruning JB. Wegener KL, Polyak SW. *Mycobacterium tuberculosis* Dethiobiotin synthetase facilitates nucleoside triphosphate promiscuity through alternate binding modes. ACS Catal. 2018 Oct;8(11):10774–10783. Available from: DOI: 10.1021/acscatal.8b03465.

21. Schumann NC, Lee KJ, Thompson AP, Salaemae W, Pederick JL, Avery T, Gaiser BI, Hodgkinson-Bean J, Booker GW, Polyak SW, Bruning JB, Wegener KL, Abell AD. Inhibition of *Mycobacterium tuberculosis* Dethiobiotin synthetase (*Mt*DTBS): toward next-generation antituberculosis agents. ACS Chem Biol. 2021 Sept;16(11):2339–2347. Available from: DOI: 10.1021/acschembio.1c00491.

22. Evans DA, Black WC. Total synthesis of (+)-A83543A [(+)-lepicidin A]. J. Am. Chem. Soc. 1993 Jun;115(11):4497–4513. Available from: DOI: 10.1021/ja00064a011.

23. Matson SL, Chatterjee M, Stock DA, John EL, Dumas EA, Ferrante CD, Monahan WE, Cook LS, Watson J, Cloutier NJ, Ferrante MA, Houston JG, Banks MN. Best practices in compound management for preserving compound integrity and accurately providing samples for assays. J Biomol Screen. 2009 Jun;14(5):476–484. Available from: DOI: 10.1177/1087057109336593.

24. Gomez-Sanchez R, Besley S, Quayle J, Green J, Warren-Godkin N, Areri I, Zeliku Z. Maintaining a High-Quality Screening Collection: The GSK Experience. SLAS Discov. 2021 Sept;26(8):1065–1070. Available from: DOI: 10.1177/24725552211017526.

25. Ludwig C, Guenther UL. Ligand based NMR methods for drug discovery. Front Biosci (Landmark Ed). 2009 Jan;14(12):4565–4574. Available from: DOI: 10.2741/3549.

26. LeBlanc RM, Mesleh MF. A drug discovery toolbox for Nuclear Magnetic Resonance (NMR) characterization of ligands and their targets. Drug Discov Today Technol. 2020 Dec;37:51–60. Available from: DOI: 10.1016/j.ddtec.2020.11.008

27. Dolomanov OV, Bourhis LJ, Gildea RJ, Howard JAK, Puschmann H. OLEX2: a complete structure solution, refinement and analysis program. J Appl Cryst. 2009 Apr;42(2):339–341. Available from: DOI: 10.1107/S0021889808042726.

28. Sheldrick GM. A short history of SHELX. Acta Crystallogr A. 2008 Jan;64(Pt 1):112–122. Available from: DOI: 10.1107/S0108767307043930.

29. Sheldrick GM. Crystal structure refinement with SHELXL. Acta Crystallogr C Struct Chem. 2015 Jan;71(Pt 1):3–8. Available from: DOI: 10.1107/S2053229614024218.

30. Gottlieb HE, Kotlyar V, Nudelman A. NMR chemical shifts of common laboratory solvents as trace impurities. J Org Chem. 1997 62(21):7512–7515. Available from: DOI: 10.1021/jo971176v.

31. Cellier M, James AL, Orenga S, Perry JD, Rasul AK, Robinson SN, Stanforth SP. Novel chromogenic aminopeptidase substrates for the detection and identification of clinically important microorganisms. Bioorg Med Chem. 2014 22(19):5249–5269. Available from: DOI: 10.1016/j.bmc.2014.08.004.

