## Supporting Tables and Figures for "Fortuitous *in vitro* compound degradation produces a tractable hit against *Mycobacterium tuberculosis* dethiobiotin synthetase: a cautionary tale of what goes in, does not always come out"

### Supporting Information

Contents


| Contents | | Page |
|---|---|---|
| Table S1 | Properties of hits identified by *in silico* screening and SPR binding analysis | 2-3 |
| Table S2 | X-Ray Crystallography Data Collection and Refinement Statistics for *Mt*DTBS complex | 4 |
| Table S3 | Crystal data and structure refinement for B9 (Compound **1**) | 5 |
| Table S4 | Key $^{1}$H NMR signals for assigning relative stereochemistry of compound **3** | 5 |
| Figure S1 | Dose-response SPR-binding analysis | 6 |
| Figure S2 | A single concentration inhibitory assay | 7 |
| Figure S3 | Comparison of 1H NMR spectra for compounds **1** and **3** | 8 |

**Table S1** Properties of hits identified by *in silico* screening and SPR binding analysis

| Compound - Structure | | ZINC ID | Affinity (kcal/mol) | MW | $K_D$ (mM) |
|---|---|---|---|---|---|
| B1 | 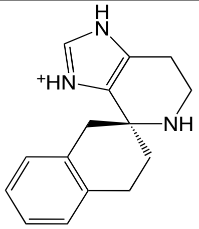 | 65409233 | -8.1 | 239.3 | 1.24 ± 0.18 |
| B2 | 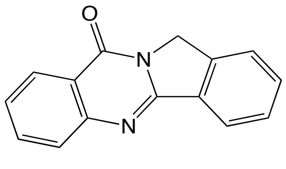 | 00487559 | -9.2 | 234.3 | ND |
| B3 | 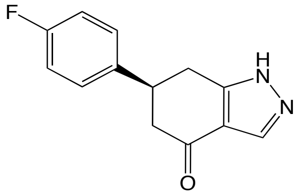 | 08771427 | -8.8 | 230.2 | 0.62 ± 0.14 |
| B4 | 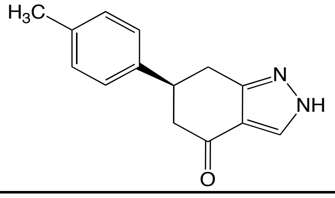 | 08738503 | -8.8 | 226.3 | ND |
| B5 | 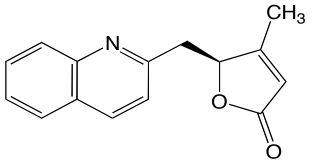 | 00262731 | -8.0 | 239.3 | ND |
| B6 | 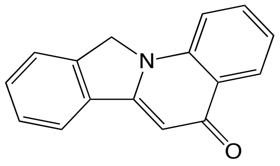 | 00519029 | -8.2 | 234.3 | ND |
| B7 | 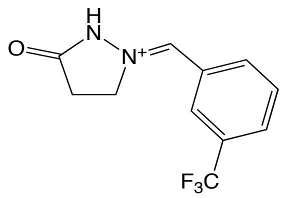 | 12956616 | -8.3 | 242.2 | 0.68 ± 0.03 |
| B8 | 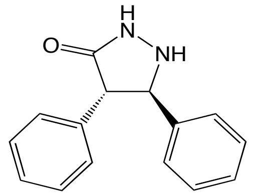 | 00083454 | -8.8 | 238.3 | ND |

ND is not detectable.

**Table S1** (continued) Properties of hits identified by *in silico* screening and SPR binding analysis

| Compound - Structure | | ZINC ID | Affinity (kcal/mol) | MW | $K_D$ (mM) |
|---|---|---|---|---|---|
| B9 | 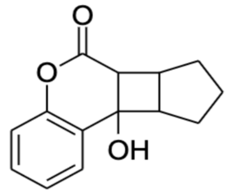 | 04114260 | -8.1 | 230.3 | 0.58 ± 0.04 |
| B10 | 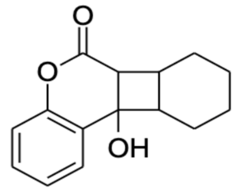 | 04114249 | -8.1 | 244.3 | ND |
| B11 | 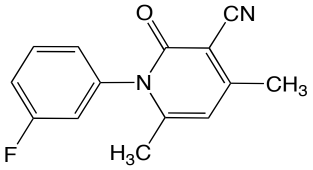 | 00054281 | -8.0 | 242.3 | ND |
| B12 | 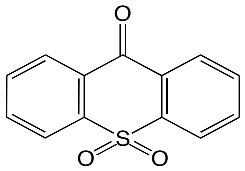 | 00400596 | -8.1 | 244.3 | ND |
| B13 | 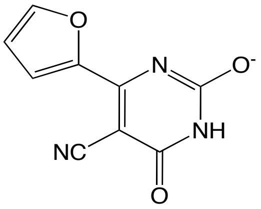 | 04958092 | -7.6 | 202.2 | ND |
| B14 | 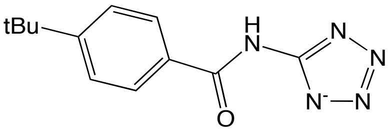 | 20149317 | -8.9 | 244.3 | ND |
| B15 | 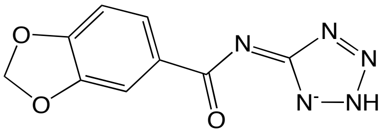 | 23382588 | -8.8 | 232.2 | ND |

ND is not detectable.

**Table S2** X-Ray Crystallography Data Collection and Refinement Statistics for the *Mt*DTBS-**3** (produced during soaking) complex. Highest shell statistics shown in parentheses.

| | ***Mt*DTBS complex with 3**<br>**PDB ID: 6NLZ** |
|---|---|
| **Wavelength** (Å) | 0.9537 |
| **Resolution range** (Å) | 62.4 - 1.90 (1.94 - 1.90) |
| **Space group** | $P2_12_12_1$ |
| **Unit cell** (Å); (°) | 54.6 105.7 154.5; 90.0 90.0 90.0 |
| **Unique reflections** | 71473 (4544) |
| **Multiplicity** | 13.0 (13.2) |
| **Completeness (%)** | 100 (100) |
| **Mean I/sigma(I)** | 14.7 (1.90) |
| **Wilson B-factor** | 22.18 |
| **R-merge (all I+ and I-)** | 0.115 (1.82) |
| **R-meas (all I+ and I-)** | 0.119 (1.89) |
| **R-pim (all I+ and I-)** | 0.033 (0.517) |
| **CC1/2** | 0.999 (0.811) |
| **Reflections used in refinement** | 70940 |
| **Reflections used for R-free** | 3490 |
| **R-work** | 0.231 |
| **R-free** | 0.273 |
| **Number of non-hydrogen atoms** | 7171 |
| **macromolecules** | 6370 |
| **ligands** | 184 |
| **solvent** | 617 |
| **Protein residues** | 909 |
| **RMS(bonds)** | 0.009 |
| **RMS(angles)** | 1.33 |
| **Ramachandran favored (%)** | 98.2 |
| **Ramachandran allowed (%)** | 1.6 |
| **Ramachandran outliers (%)** | 0.22 |
| **Rotamer outliers (%)** | 1.6 |
| **Clashscore** | 6.9 |
| **Average B-factor** ($Å^2$) | 40.4 |
| **macromolecules** ($Å^2$) | 40.6 |
| **ligands** ($Å^2$) | 32.2 |
| **solvent** ($Å^2$) | 41.6 |

**Table S3** Crystal data and structure refinement for B9 (Compound 1)

| | |
|---|---|
| Identification code | B9/Compound **1** |
| Empirical formula | $C_{28}H_{28}O_6$ |
| Formula weight | 460.50 |
| Temperature/K | 150(2) |
| Crystal system | triclinic |
| Space group | *P*-1 |
| a/Å | 8.3370(5) |
| b/Å | 10.4025(7) |
| c/Å | 13.4821(5) |
| α/° | 106.663(5) |
| β/° | 90.064(4) |
| γ/° | 103.109(5) |
| Volume/Å$^3$ | 1088.21(11) |
| Z | 2 |
| $\rho_{calc}$g/cm$^3$ | 1.405 |
| μ/mm$^{-1}$ | 0.098 |
| F(000) | 488.0 |
| Crystal size/mm$^3$ | 0.95 × 0.59 × 0.21 |
| Radiation | Mo Kα (λ = 0.71073) |
| 2Θ range for data collection/° | 6.97 to 58.564 |
| Index ranges | -11 ≤ h ≤ 11, -13 ≤ k ≤ 13, -18 ≤ l ≤ 18 |
| Reflections collected | 19114 |
| Independent reflections | 5199 [$R_{int}$ = 0.0382, $R_{sigma}$ = 0.0367] |
| Data/restraints/parameters | 5199/0/309 |
| Goodness-of-fit on $F^2$ | 1.030 |
| Final R indexes [I>=2σ (I)] | $R_1$ = 0.0469, $wR_2$ = 0.1126 |
| Final R indexes [all data] | $R_1$ = 0.0605, $wR_2$ = 0.1219 |
| Largest diff. peak/hole / e Å$^{-3}$ | 0.38/-0.26 |

**Table S4** Key $^1$H NMR signals for assigning relative stereochemistry of compound **3**

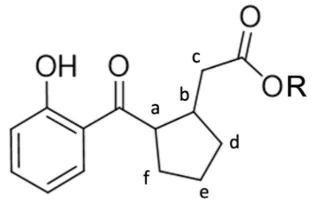


**3** R= H

**4** R= $CH_2CH_3$

| | 1H Chemical shift (ppm) | | |
|---|---|---|---|
| **$^1$H** | **Compound 3** | **Cis-4** | **Trans-4** |
| **a** | 3.52 | 4.06 – 3.87* | 3.58 – 3.51 |
| **b** | 2.89 | 2.83 – 2.73 | 2.95 – 2.83 |
| **c** | 2.47, 2.37 | 2.34 – 2.22 | 2.45 – 2.33 |
| **d/e/f** | 2.23 – 2.06, 1.83 – 1.71, 1.51 – 1.37 | 2.13 – 2.04, 2.03 – 1.84, 1.73 – 1.59 | 2.21 – 2.01, 1.86 – 1.69, 1.48 – 1.38 |

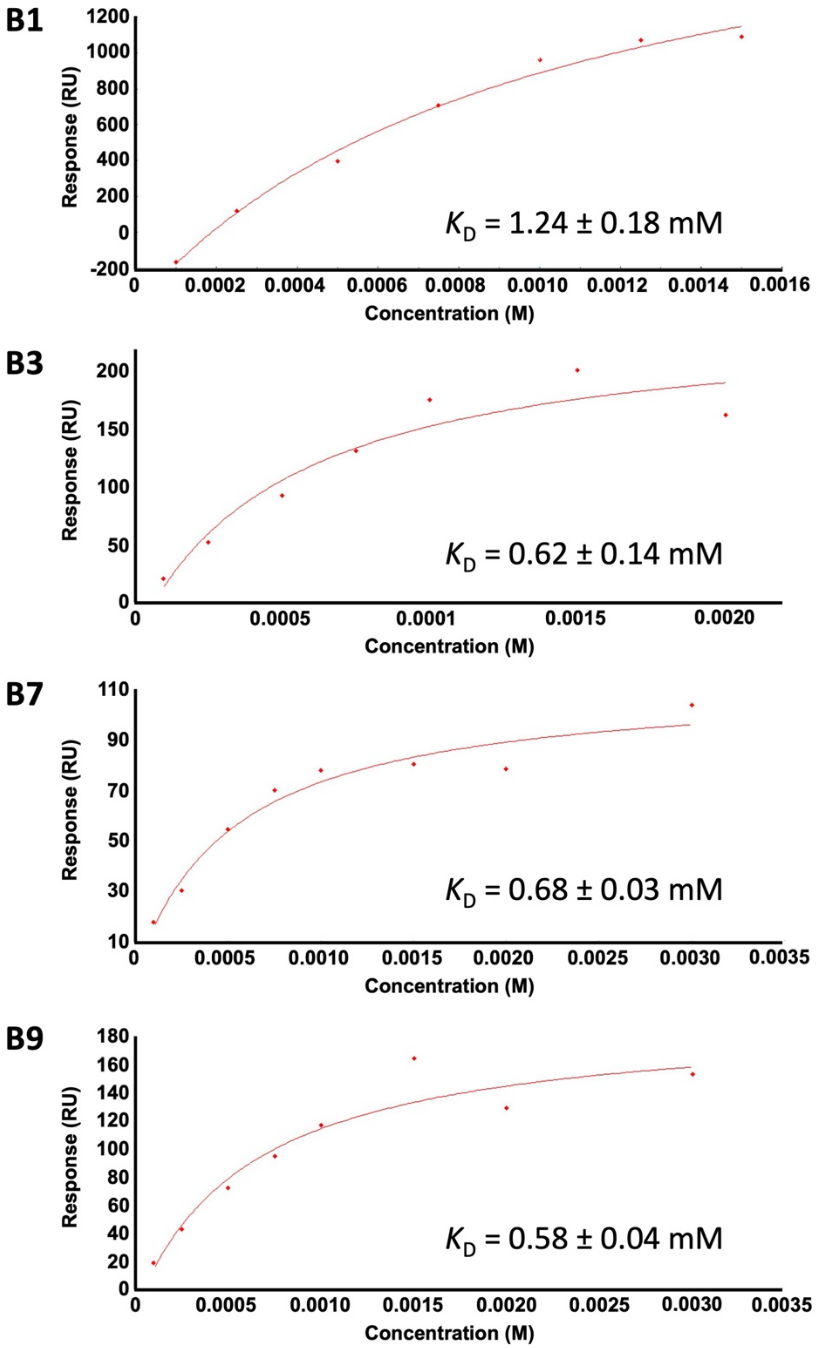


**Figure S1** Dose-response SPR-binding analysis. Binding analysis based on affinity-steady state 1:1 model. The dose response curves of **B1**, **B3**, **B7**, and **B9** represent a single experiment. The binding affinity ($K_D$) of fragment to *Mt*DTBS was calculated from three independent experiments (n=3).

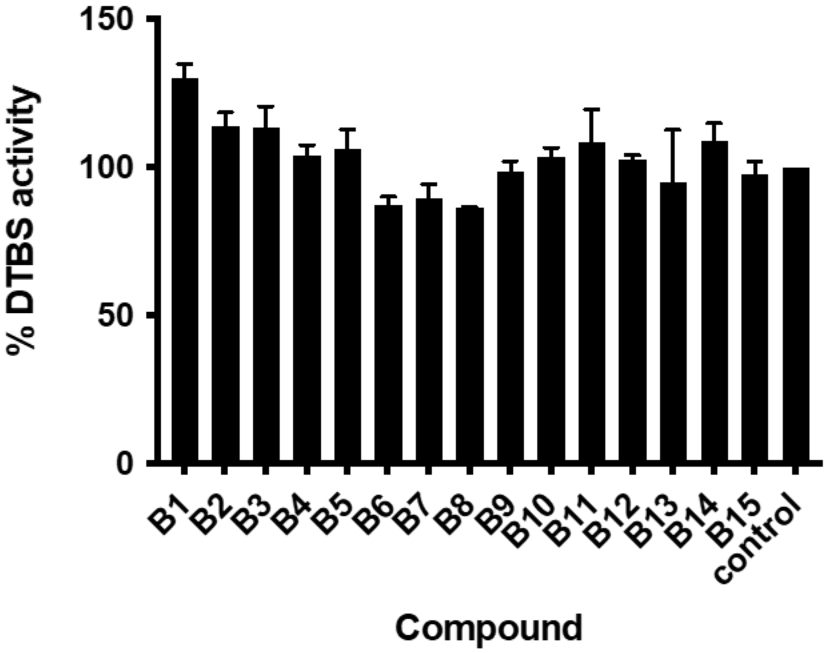


**Figure S2** A single concentration inhibition assay. The inhibitory activity of 1% DMSO-solubilized compounds against *Mt*DTBS is presented as the percentage of enzyme activity. Reaction without compound was normalised to 100% (shown as the control bar). The test concentration (mM) of compounds (B1, 0.7; B2, 0.27; B3, 1.45; B4, 1.77; B5, 1.67; B6, 0.18; B7, 1.38; B8, 1.05; B9, 2.39; B10, 2.07; B11, 0.27; B12, 0.28; B13, 2.15; B14, 0.50; B15, 2.11) was the maximum concentration in which each compound could be solubilized in the 1% DMSO assay. The data represents mean ± SD of three independent experiments.

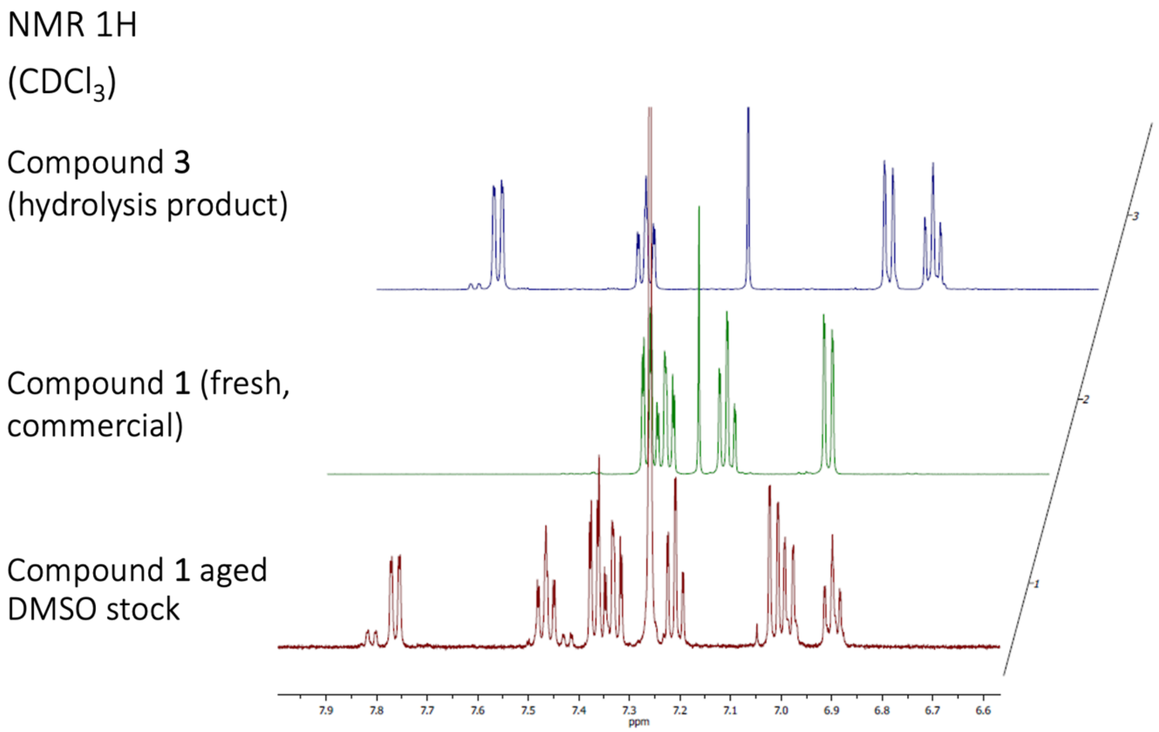


**Figure S3** Comparison of 1H NMR spectra of aged DMSO stock of compound **1** (red), to freshly obtained commercial **1** (green) and intentionally produced compound **3** (blue). The DMSO stock contained a mixture of **1** and **3.**
